# Regionalization in the compound eye of *Talitrus saltator* (Crustacea, Amphipoda)

**DOI:** 10.1101/844027

**Authors:** Alice Ciofini, Luca Mercatelli, Yumi Yamahama, Takahiko Hariyama, Alberto Ugolini

## Abstract

The crustacean *Talitrus saltator* is known to use many celestial cues during its orientation along the sea-land axis of sandy shores. In this paper, we investigated the existence of the eye regionalization by morphological, electrophysiological and behavioural experiments. Each ommatidium possesses five radially arranged retinular cells producing a square fused rhabdom by R1-R4 cells; the smaller R5 exist between R1 and R4. The size of R5 rhabdomere is largest in dorsal part and becomes gradually smaller in median and ventral part of the eye. Spectral-sensitivity measurements were recorded from either dorsal or ventral parts of the compound eye to clarify the chromatic difference. Results show that the dorsal part is green and UV-blue dichromatic, whereas the ventral part is UV (390 nm) with a substantial population of 450 nm receptors with the responses in the longer wavelength region. To evaluate the orienting behaviour of individuals, their eyes were black painted either in the dorsal or ventral part, under natural sky or a blue filter with or without the vision of the sun. Results show that animals painted on the dorsal part of their eyes tested under the screened sun were more dispersed and in certain cases their directions deflected than other groups of individuals. Furthermore, sandhoppers subjected to the obscuring of this area met in any case high difficulties in their directional choices. Therefore, our present work indicates the existence of a regionalization of the compound eye of *T. saltator*.

**Summary statement:** This work provides evidences of the morphological and electrophysiological regionalization of the compound eye and the visual capabilities for behaviour involved in the recognition of the celestial compass orienting factors in crustaceans.

## INTRODUCTION

The supralittoral sandhopper *Talitrus saltator* (Montagu, 1808) to return as quick as possible to the belt of damp sand in which it lives buried during the day orientates along the sea-land axis of the beach, perpendicular to the shoreline (characteristic for each population dependently on the orientation of the shore). This species to avoid high temperature and low humidity conditions, which can rapidly lead animals to death, evolved nocturnal habits. However, stressing factors, such as predators, strong winds or changes in the sea level, can cause active displacements of sandhoppers from their preferred zone also during the day. In any case, animals orientate seaward when exposed to dry conditions whereas submersion motivates them to direct landward (Pardi and Papi, 1952).

In its zonal recovery, *T. saltator* is known to rely on several celestial cues; of course, their use requires continuous adjustments to account for their temporal azimuthal variations and maintain a constant effective direction. Evidence for time-compensated solar and lunar orientation has been provided since the fifties (see Pardi and Ercolini, 1986 for a review) and recently the skylight intensity gradient has been found to provide reliable directional information to this species (Ugolini et al., 2009). Furthermore, the solar orientation of *T. saltator* depend on a skylight cue perceived in the UV-blue band whose nature has not been identified yet. Indeed, when the perception of wavelengths shorter than 450 nm is prevented sandhoppers lose their capability to recognize the sun as a compass cue and exhibit positive phototactic tendencies (Ugolini et al., 1993, 1996). The sensitivity of sandhoppers to UV-blue light has recently clearly confirmed by electrophysiological recordings (Ugolini et al., 2010). In fact, it has been revealed the existence of discrete photoreceptors within the compound eye of *T. saltator* sensitive to UV-blue (λ = 390-450 nm) and green (λ = 500-550 nm) wavelengths, respectively.

The aim of this work is to assess the eventual regionalization of the visual capabilities involved in celestial compass orientation in the sandhopper *T. saltator* through morphological, electrophysiological and behavioural investigations.

## METHODS

Adult individuals of *T. saltator* were collected on a sandy beach in the Regional Natural Park of Migliarino, San Rossore, Massaciuccoli, Pisa, Italy (43°40’03”N, 10°20’29”E, sea-land axis = 265°-85°) over spring-summer 2016. After the collection, animals were transported to the laboratory of University of Florence and kept in Plexiglas boxes containing damp sand; food (universal dried fish food, SERA, Vipan, Heisemberg, Germany) was available *ad libitum*. Sandhoppers were maintained at ambient temperature (25°C) under an L:D = 12:12 cycle in phase with the natural photoperiod (sunrise = 0600 hours, sunset =1800 hours) and the behavioural experiments were performed within 14 days from their capture. All the tests were carried out with intact individuals and with individuals with the dorsal (1/3) or ventral (2/3) part of their eyes painted by black enamel (Fig. 1C, D). For the morphological and electrophysiological experiments, animals were transferred by airplane to Japan, and examined at Hamamatsu University School of Medicine.

**Fig 1.**
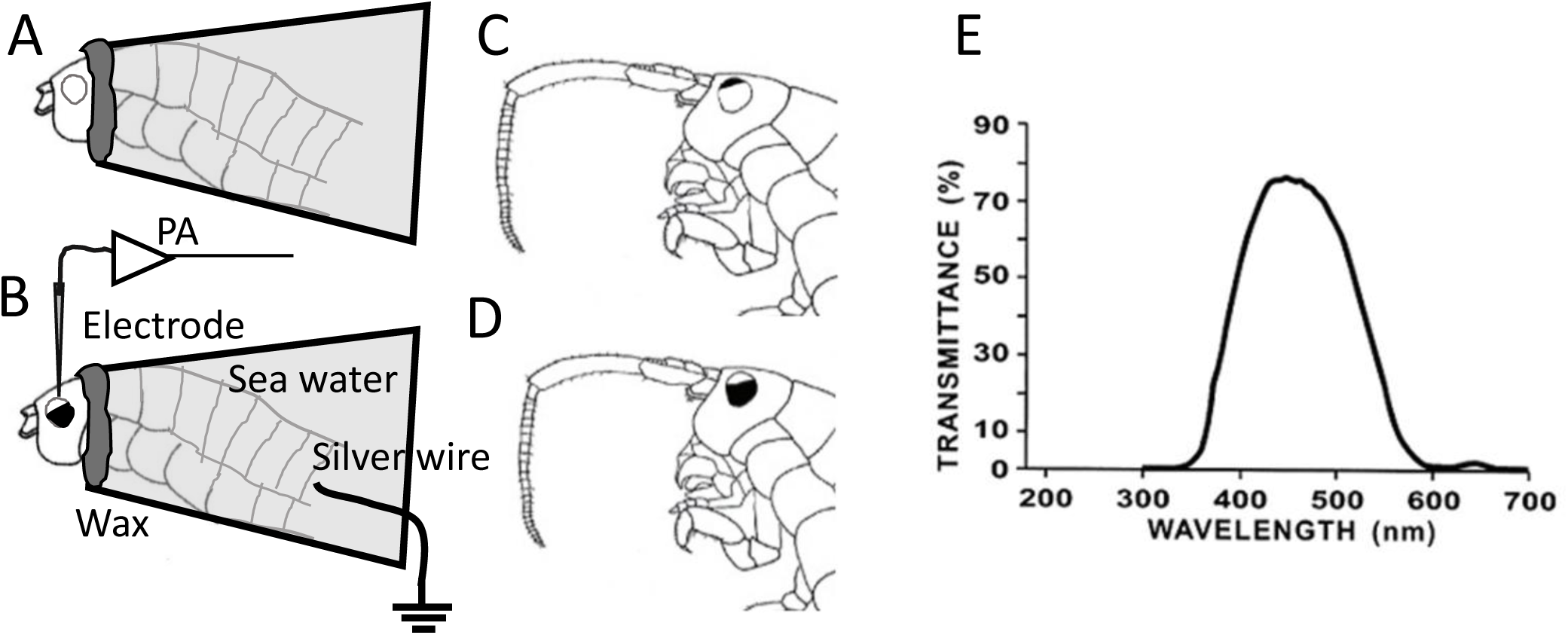
Schematic representations of the holding method for electrophysiological recording. (A) The animal cut the tail region was introduced into the plastic tube and the head was glued by beeswax and resin to separate electrically. (B) A part of the compound eye was pained and the tip of the electrode was placed just below the cornea at the centre of unpainted area. (PA) preamplifier. (C, D) sandhoppers with the dorsal 1/3 region or 2/3 ventral region of their eyes painted, respectively. (E) transmittance curve of the Blue gelatine filter.

### Morphological investigations by microscopy

#### SEM

NanoSuit method was used to observe *T. saltator* (Ohata et al., 2014). Briefly, the whole body of living sandhoppers were dipped into the NanoSuit solution which contains mainly 1% (volume percent) polyoxyethylene (20) sorbitan monolaurate solution dissolved in distilled water for ca.10 sec, blotted briefly on a dry filter paper to remove excess solution. Then, the living specimen was introduced into the SEM to construct a NanoSuit, a very thin film on the surface of the specimen, by the irradiation of electron beams, without performing any traditional treatments such as chemical fixation or dehydration. The head of the living sandhopper was observed by FE-SEM (Hitachi S-4800) at an acceleration voltage of 5.0 kV.

#### TEM

The compound eye is easily broken by the pressure from scissors or a scalpel when attempting to remove the eye from the body. Therefore, the head was removed gently from the body in a primary fixative solution (2% glutaraldehyde with 2% paraformaldehyde in 0.1 mol l^−1^ sodium cacodylate buffer, pH 7.2) with a small piece of razor blade, and kept for 2 hours at 4°C. The fixed heads were dissected through the median line of the head in a buffered solution, and were then postfixed for 1 hour in 1% OsO_4_ in 0.1 mol l^−1^ cacodylate buffer and washed three times for 10 min each in 0.1 mol l^−1^ cacodylate buffer. Dehydration through a graded series of ethanol solutions was followed by embedding in an Epon–Araldite mixture. Ultrathin (approximately 70 nm) sections were cut perpendicular or parallel to the optical axis of each ommatidium and stained with 2% uranyl acetate followed by 0.4 % lead citrate for 5 min each. Observations were performed with JEOL JEM-1400 transmission electron microscope (100kV).

### Measurement of the visual field of the unpainted area

To measure the vertical visual filed of the unpainted area both of the DP (painted the 2/3 ventral part of the compound eye, Fig. 1D) and VP (painted the 1/3 dorsal part, Fig. 1C), we made light microscopic observations of the semi-thin section of the eye at the centre of the compound eye in the transverse plane. The embedded specimen in an Epon–Araldite mixture same as the TEM observation were sectioned parallel to the optical axis of each ommatidium and stained with Toluidine blue. We lined the optical axis between the centre of the proximal region and the distal end of the crystalline cone at the both ends of the unpainted area (Fig.3 A, D) to find the line of the dichotomy between the dorsal and ventral visual field predicted.

### Electrophysiological recordings

Because of the easy deformation of the eye structure when the body is pressed, the animal was gently treated not to give any mechanical stress. Therefore, the whole animal was used but only the tail region was cut in the seawater to prevent from the mechanical stress. The whole body without tail region was put into the plastic tube, and placed only the head part including the compound eyes out of the tube. The body side of the head was glued to a tube using melted beeswax and resin (1:1) (Fig. 1A). The body was placed inside of the tube, and was filled with artificial seawater. The indifferent electrode, a chloride silver wire, was placed into the tube (Fig. 1B) to connect electrically through the cut area at the tail.

To compare the spectral response curve measured between dorsal part (DP) and ventral part (VP), we prepared the animal painting with black enamel (Touch up paint T-13, SOFT99 Corp.) the dorsal or ventral parts of the eyes. To distinguish the response according with the region of DP and VP, the recording electrode was placed around the centre of unpainted area. a small hole (ca. 50 µm, crystalline cone diameter) was made on the surface of the cornea using a razor blade, and the tip of the glass electrode filled with artificial seawater was inserted to the hole just on the surface of the crystalline cone layer for electroretinogram (ERG) recordings. The end of the light guide was placed near the compound eye of unpainted area. With this location of electrodes, we obtained stable cornea negative responses. We also excluded the responses of on- and off-transients. Because transient responses are the pooled activity of second order neurons of the lamina, whereas the sustained photoreceptor response is the pooled activity of photoreceptors and is easily modified by transient responses (Heisenberg, 1971).

Responses were amplified with a high-impedance preamplifier (Nihon Kohden MZ 8201) and a high-gain amplifier (Nihon Kohden AVH-10). The magnitude of the responses (peak amplitude) was monitored on an oscilloscope (Nihon Kohden VC10). Permanent recordings were made with a data logger (Keyence Corp. NR-600). An optical system was used, with a 500-W Xenon arc lamp (Ushio Inc., Type UXL-500D-O, Japan) with a regulated power supply (Sanso XD-25, Japan). Quartz lenses produced a parallel beam of light which passed through a heat-absorption filter (Toshiba IRA-25S), and a set of 17 narrow-band interference colour filters (Vacuum Optics Corp., Japan) scanning a range of 330 to 650 nm having half-bandwidth maxima of 15 nm. The light intensities of the testing were measured with a silicon photodiode (S876-1010BQ, Hamamatsu Photonics K.K.) calibrated by Hamamatsu Photonics K.K. using a photo-electron bulb, and each monochromatic light was adjusted with the aid of several neutral density filters so that each contained an equal number of photons. At all wavelengths, the maximum irradiances available at the surface of the eye were 1.3 × 10^14^ quanta/ cm^2^/s. The light intensity was altered with quartz neutral-density filters (Vacuum Optics Corp., Japan) to obtain intensity-response (V-log I) curves. To get the spectral-response curves, the light irradiance of each testing light was 1.3 × 10^13^ quanta/cm^2^/s, which corresponds with 1 log unit lower than the maximum irradiance. Each beam was interrupted by shutters (Uniblitz photographic and Copal) controlled for the duration of the test flash and adaptation times by a stimulator (Nihon Kohden SEN-7103). The eye was stimulated by a 200-ms pulse of light, and the stimulus interval was 20 s. Experiments were performed from 12:00 to 17:00 h during the day.

### Celestial compass experiments

Behavioural experiments were carried out in a confined environment by using a device similar to that described by Ugolini and Macchi (1988). It consisted on a tripod supporting a horizontal transparent plate (diameter = 30 cm) with a Plexiglas bowl (diameter = 20 cm) placed on where sandhoppers were released. A goniometer was set under the bowl to detect the directions assumed by individuals. The vision of the surrounding landscape was prevented by using an opaline Plexiglas screen (diameter = 30 cm, height = 3 cm).

Sandhoppers were dehydrated for a few minutes before being tested in order to motivate them to orientate seaward. Groups of 10-15 individuals were released into the bowl at a time since it has been demonstrated their non-reciprocal influence in performing directional choices (Scapini et al., 1981). A single direction (error ± 2.5°) for each radially-oriented animal (with the head pointed toward the outside of the bowl and the longitudinal axis of the body oriented no more than **±** 45° from the radius of the bowl) was measured from freeze-frame images recorded by using a video-camera placed under the apparatus. In each trial, even the number of radially orientated individuals out of the total number of individuals released was registered.

In order to assess the eventual regionalization of the visual pigments within the compound eye of this species, we tested separately both intact individuals and sandhoppers subjected to the occlusion of discrete regions of their eyes. In particular, we performed the following experimental treatments:

1. painting of the dorsal part (DP, corresponding to the 1/3 upper area) of their eyes with black enamel (Rainbow, Maimeri, S.p.A., Mediglia, Milano, Italy) (Fig. 1C).
2. painting of their whole eyes except for the dorsal edge (the 2/3 ventral part, VP) (Fig. 1D).

Releases were carried out both under the natural sky and with the superimposition of a gelatine blue filter (Moonlight Blue, no 183, Spotlight, Milano, Italy, λ_max_ = 450 nm, transmittance = 73%, Fig. 1E).

Tests were repeated in conditions of sun visible and with its vision prevented by using a screen (42 × 42 cm) placed at a distance of about 2 m from the bowl.

Data were analysed according to the methods proposed by Batschelet (1981) for circular distributions.

The length of the mean resultant vector (*r*) and the mean angle were calculated (α). To establish whether the distributions differed statistically from uniformity the *V*-test was used (*P*<0.05 at least). For each distribution the goodness of orientation (*v* = *r* cos |α -δ|, δ = expected direction, ranging from 0 to 1) was measured to estimate the concentration of the directional choices exhibited by the individuals released and the deflection of the mean resultant vector with respect the expected direction.

Furthermore, in order to evaluate if discrete distributions were statistically different from each other we carried out pairwise comparisons by using the Watson’s *U*^*2*^ test (*P*<0.05 at least).

Since the frequency of radially-orientated individuals is considered a good indicator to assess the difficulty of sandhoppers in their directional choices we conducted statistical comparisons between frequencies recorded in different trials by using the *G*-test (*P*<0.05 at least) (Zar, 1984).

## RESULTS

### Morphological investigations by microscopy

#### Optical and SEM

The compound eye of *T. saltator* is sessile, and each ommatidium is observed as black dots using a light microscope (Fig.2 A). The cornea of each ommatidium possesses no curvature and entire cornea of the compound eye also has no curvature. The surface of the cornea covering the eye is smooth and not divided into facets as in many other compound eyes when observed by the NanoSuit method using a SEM (Fig.2 B). TEM observation revealed that each ommatidium is located beneath an undifferentiated cuticlar lens and the existence of the long crystalline cone. The length of the crystalline cone occupied half of the ommatidium (Fig.2 C). Following the crystalline cone, there are rhabdoms produced by retinular cells without crystalline tract, therefore the compound eye of *T. saltator* is classified in apposition-type.

**Fig. 2.**
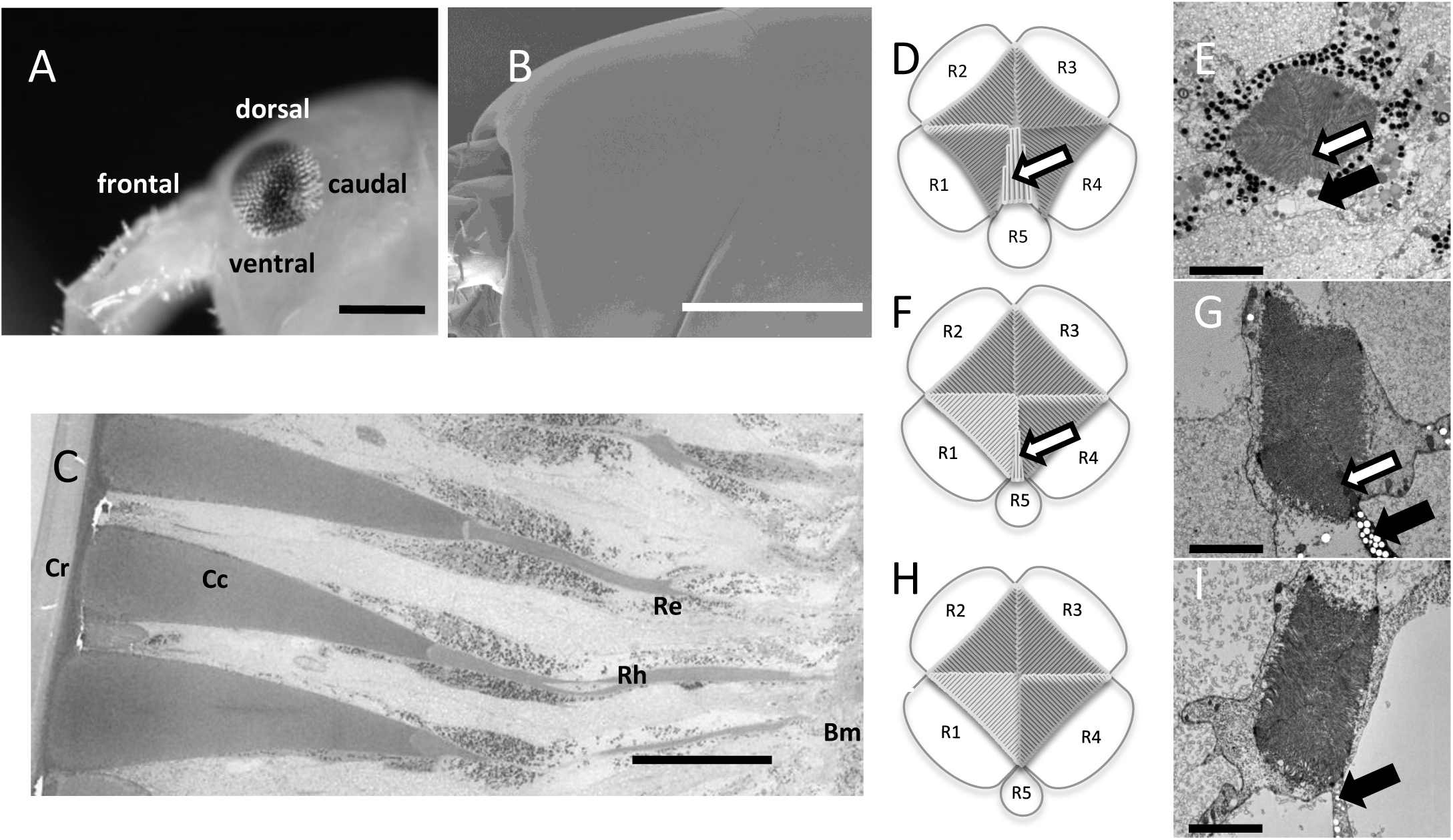
Morphological characterization of the compound eye. (A) A light microscopic view of the head shows each ommatidium through the transparent cornea. (B) Scanning electron microscope of the region of compound eye observed by the NanoSuit method shows the smooth surface of cornea and not divided into facets as in many other compound eyes. (C) Transmission electron micrograph of the longitudinal section of three ommatidia. The scale bars in A and B indicate 500 μm. The scale bar in D indicate 50 μm. The schematic drawings (D, F, H) and the transmission electron micrograph of the rhabdom (E, G, I), obtained from the dorsal region (D, E), median region (around the equatorial line) (F, G) and the ventral region (H, I) of the compound eye. White arrows indicate each rhabdomere of R5, and Black arrows indicate the soma of R5. Scale bars indicate 5 μm. Cr: cornea, Cc: crystalline cone, Rh: rhabdome, Re: retinular cell, Bm: basement membrane.

#### TEM

The cross section of the rhabdom shows the radially arranged five retinular cells (R1-R5) producing a square rhabdom, in the centre of the retinular cells in each ommatidium. It is obvious that the square rhabdom is produce by four triangular rhabdomeres of each retinular cell (R1-4). The sizes of them are almost equal, and the microvillar orientation of the pair R1 and R3 is orthogonal to the pair R2 and R4 (Fig. 2 D-I). The smaller R5 exist between R1 and R4, and the microvillar orientation of the rhabdomere is parallel to the dorsoventral line of the compound eye. The area ratio of the R5 rhabdomere (Fig.2 D - G, white arrows) to the whole rhabdomeres (R1-4 and R5) was 6.90 ± 0.14% (n=5) in the dorsal part (Fig. 2. D, E), and 1.94 ± 0.41% (n=5) in the median part (Fig. 2. F, G). In the ventral part, the areal ratio was decreased to 0.40 ± 0.28% (n=5) even the soma of the R5 exist (Fig. 2 H, I, black arrows).

The schematic drawings of Fig. 3 A-E show the visual fields of DP or VP when they were covered by black paint. Each Θ of DP was 55°, therefore the animal might be able to obtain a visual field of ca. 90 ° by using both eyes. In contrast, the visual field of each VP was ca. 90°. Therefore, we painted the eye of 1/3 dorsal part (DP) or 2/3 ventral part (VP) by black paint in electrophysiological and behavioural experiments.

**Fig. 3.**
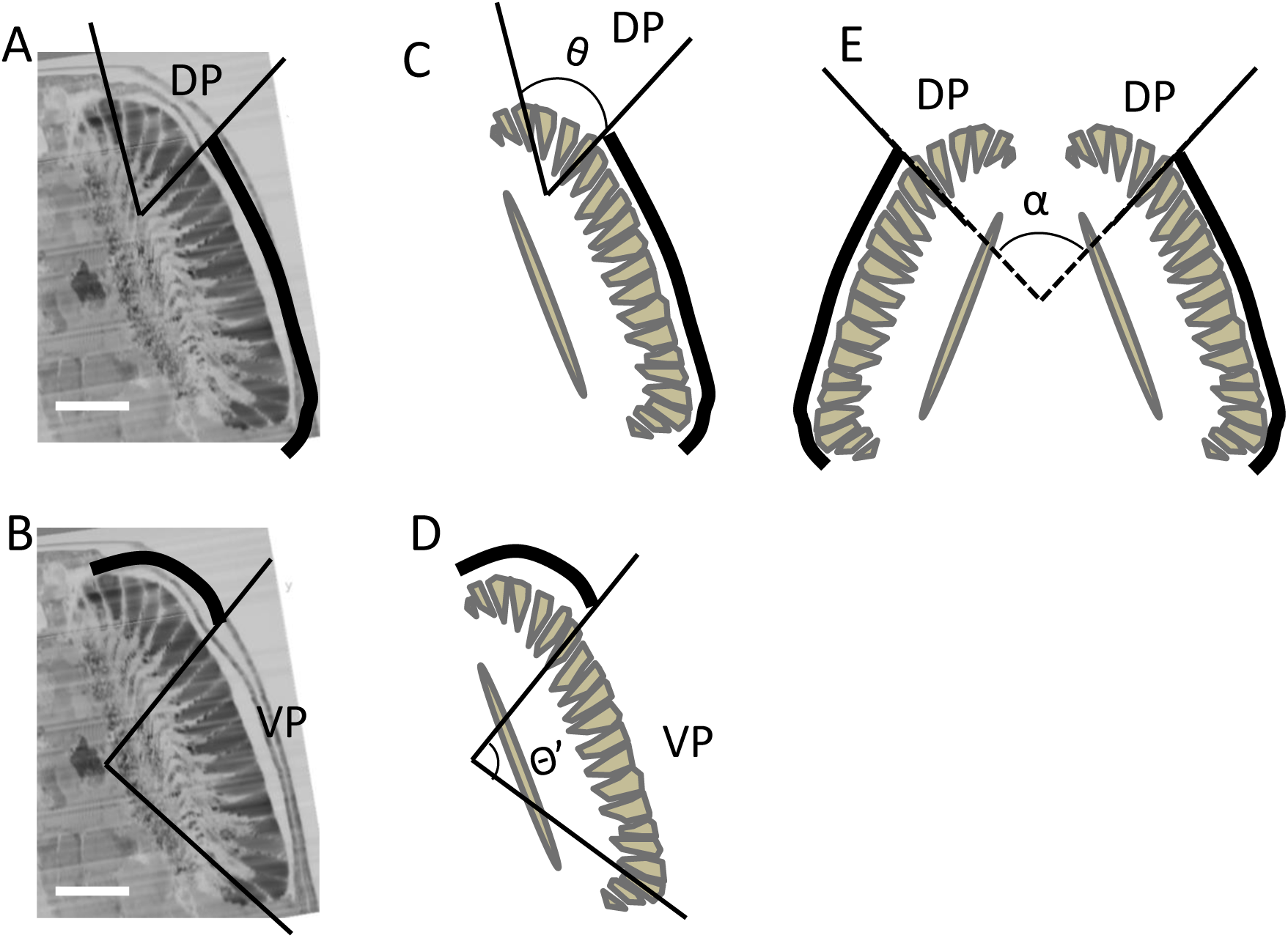
Visual field in dorsal and ventral regions. (A) and (D) light microscopic view of the longitudinal sections of the compound eye along the transverse plane. Scale bars indicate 100 *μ*m. (C, E)unpainted dorsal part. (D) unpainted ventral part. Small crystalline cones were omitted. Dorsal part Θ=60°; α=85°; Ventral part; Θ’=85°

### Electrophysiological recordings

Both the compound eye painted at the DP or VP showed the corneal-positive sustained waveform when irradiated by under around 1.0 × 10^13^ quanta/ cm^2^/s, however, on-transient waveforms were observed above the light intensity at the wavelength of peak sensitivities such as UV-blue or green region. Therefore, we stimulated by the light intensities which only caused sustained waveforms.

The relative response (V/Vmax) curves (V-log I curves) and the relative spectral sensitivity curves derived from ERG recordings are shown in Fig. 4. There was no apparent difference between dorsal and ventral parts in V-log I curves (Fig. 4 A). In the DP of the compound eye, the spectral sensitivity curves showed two distinct maxima, one in the long-wave range peaking at 550 nm and one in the UV-blue range peaking at 430 nm, divided by a sensitivity minimum at approximately 470 nm (Fig. 4 B). The spectral sensitivities of VP differed significantly from those of DP. The spectral sensitivity curves of VP possessed high sensitive region only around UV-blue peaking at 390 and 450 nm, and was decreased gradually in longer wavelength; no apparent peaking region was observed.

**Fig. 4.**
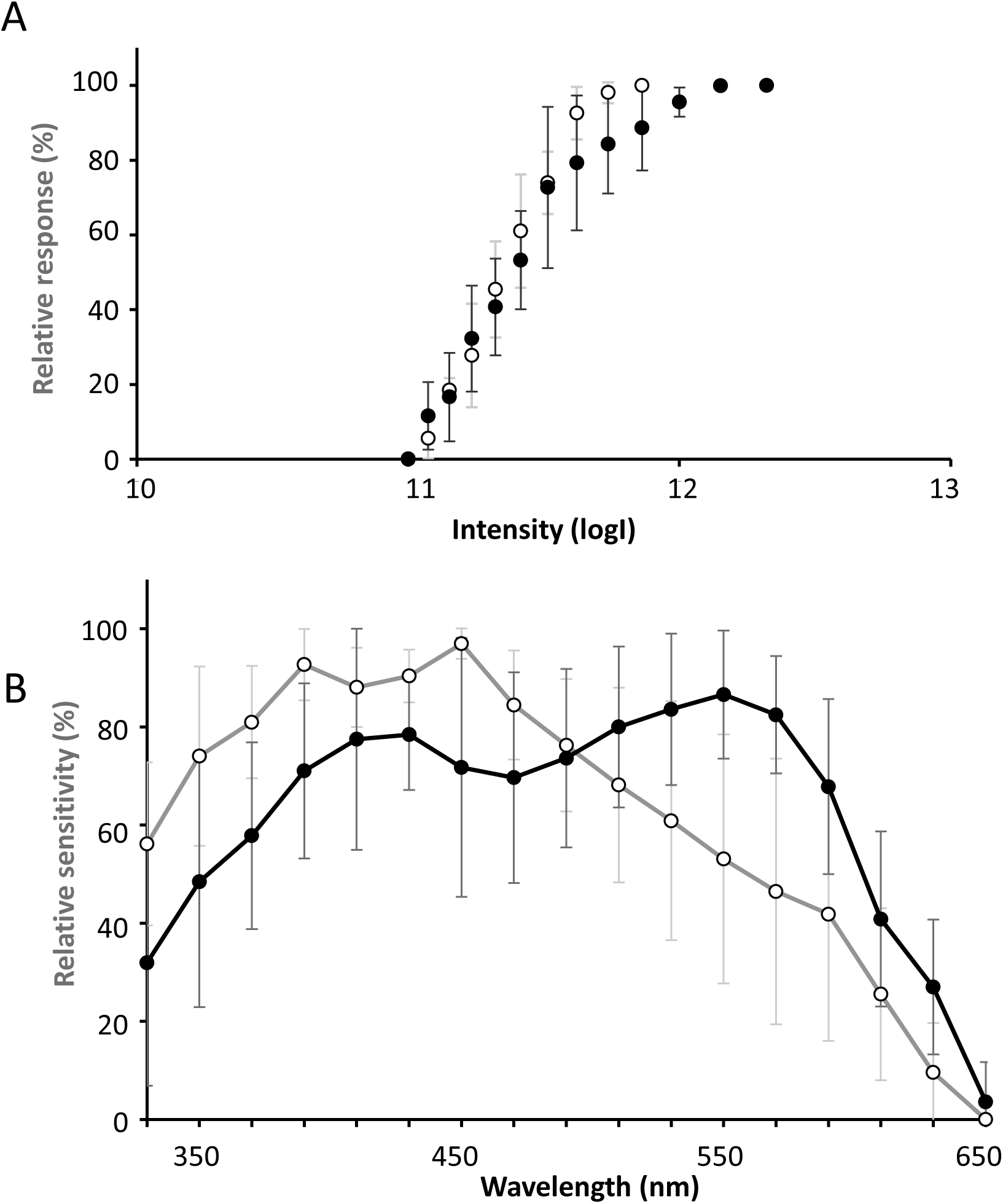
Electrophysiological recordings. (A) The V/log I function obtained for the dorsal (black dots) and ventral (white dot) part of the compound eye plotted from responses to 200 ms flashes of varying intensity of white light. Data are means (±SD, N=5). (B) Normalized, averaged (± SD, N=5) spectral sensitivity curves from dorsal (black dots) and ventral (white dots).

### Celestial compass experiments

In the releases conducted under the natural sun and sky not treated individuals (controls) exhibited a mean orientation in good agreement with the seaward direction of their home beach (*v* = 0.65, Fig. 5 A). Similarly, sandhoppers with either the dorsal part (DP) (Fig. 5 B) or the 2/3 VP (Fig.5 C) of their eyes obscured performed directional choices mainly directed toward the expected direction (*v* = 0.65 and 0.66 respectively)

**Fig. 5.**
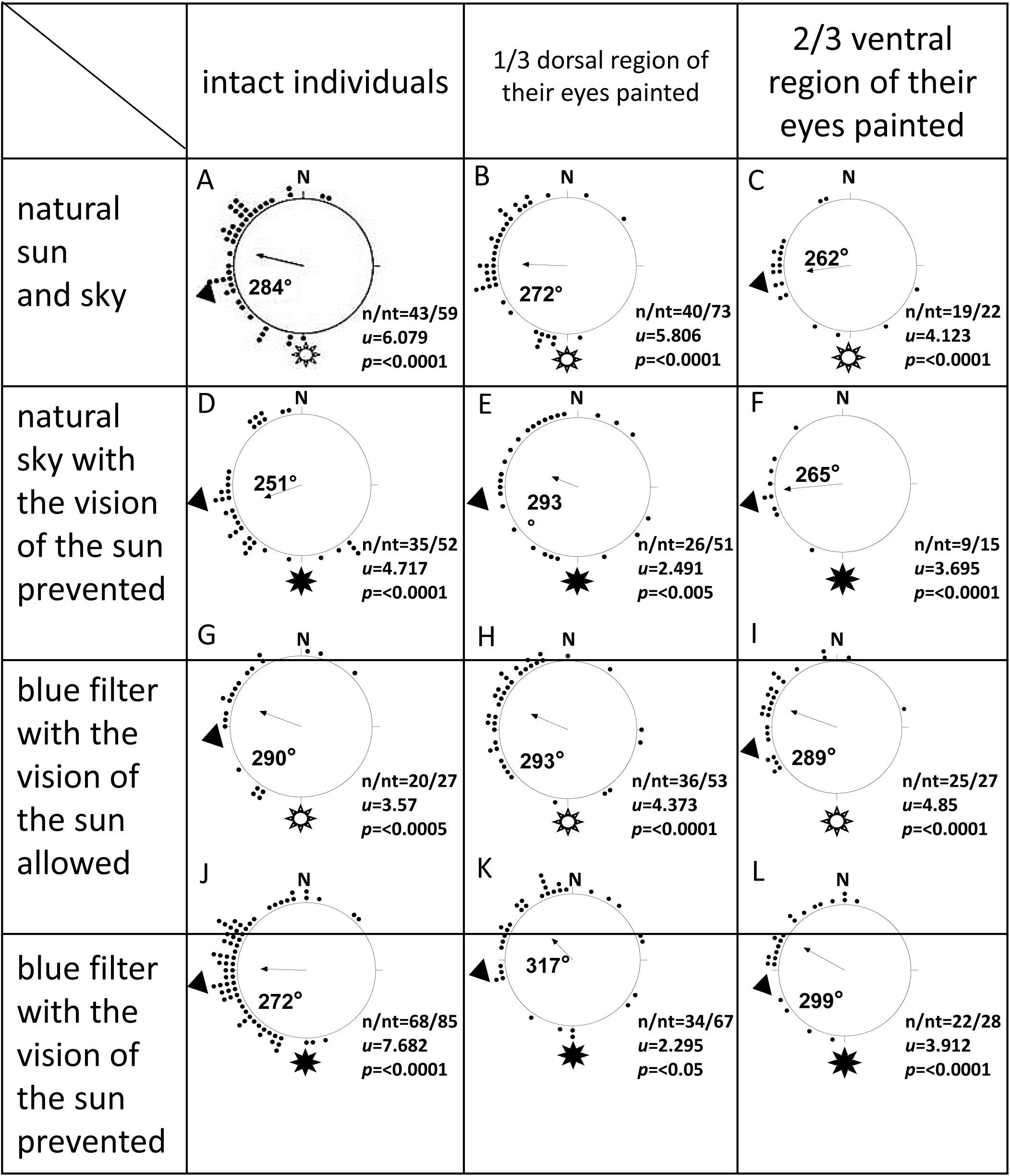
Releases under the natural sun and sky. (A) intact individuals, (B) individuals with the 1/3 dorsal region of their eyes painted, (C) individuals with the 2/3 ventral region of their eyes painted and (D) frequencies of radially-orientated sandhoppers recorded in discrete trials. N, North, black arrows, mean vector and angle (the length of the mean vector ranges from 0 to 1 = radius of the circle), black dots, sandhoppers’ directions (each dot corresponds to the direction of one individual); black triangles outside the distributions, seaward direction of animals’ home beach. The symbol of the sun indicates the solar azimuth at the time of releases. n/nt, number of radially-orientated sandhoppers (n) out of the total individuals tested (nt). The values of the V-test, u, with their probability level, P, are also given.

Pairwise comparisons between distributions conducted by the Watson’s *U*^*2*^ test did not reach the statistical significance in any case. Instead, the frequencies of radially-orientated individuals differed statistically from each other (*G*-test, *G* = 9.745, df = 2, *P*<0.01, Fig. 6 A).

**Fig. 6.**
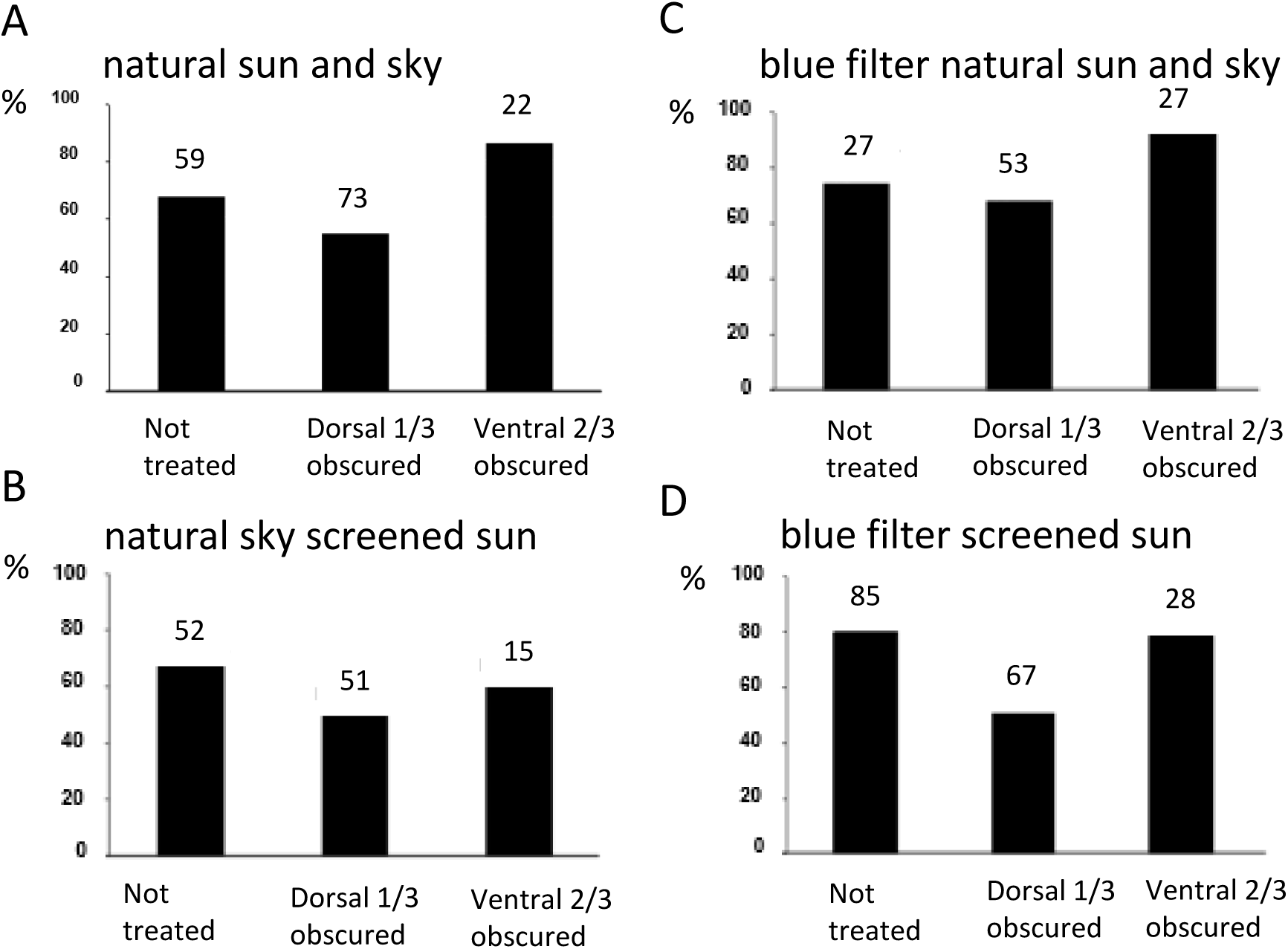
Frequencies of radially orientated sandhoppers. (A) natural sun and sky, (B) natural sky with the vision of the sun prevented, (C) blue filter with the vision of the sun allowed and (D) blue filter with the vision of the sun prevented. The total number of individuals released is given on the top of each bar. See text for further explanations.

When tested under the natural sky and with the sun screened controls individuals (Fig. 5 D) and sandhoppers with the VP of their eyes painted (Fig. 5 F) orientated in accordance with the seaward direction of their home beach (*v* = 0.56 and 0.87 respectively). Even animals with the DP exhibited a mean orientation in accordance with the expected direction (Fig. 5 E) but they were more dispersed than the other groups of individuals and the value of *v* measured in these trials (0.34) was lower than those recorded in releases of the other lots. Comparisons between distributions revealed that the distribution obtained from releases of sandhoppers with the DP of their eyes painted differed statistically from those referring to experiments conducted with intact animals (Fig.5 A *vs* B, Watson’s *U*^*2*^ test, *U*^*2*^_(43, 40)_ = 0.201, *P*<0.05) and individuals with the 2/3 ventral area of their eyes obscured (Fig. 5 B *vs* C, Watson’s *U*^*2*^ test, *U*^*2*^_(40, 19)_ = 0.205, *P*<0.05). In this case, the difference in the frequencies of radially-orientated sandhoppers registered in discrete trials did not reach the statistical significance (*G*-test, *G* = 2.778. df = 2, *P* = NS, Fig. 6 B).

In experiments carried out under the blue gelatine filter and the vision of the sun allowed, controls exhibited a mean orientation significantly concordant with the expect direction (Fig.5 G); indeed, the value of the goodness of orientation recorded was 0.56. Even the directional choices performed by individuals subjected to the occlusion of either the DP (Fig. 5 H) or the VP (Fig. 5I) of their eyes were mainly orientated in accordance with the seaward direction of their home beach; the values of *v* calculated from these trials were 0.53 and 0.68 respectively.

Pairwise comparisons carried out by the Watson’s *U*^*2*^ test revealed that the distributions did not differed statistically each other. Instead, significant differences in the frequencies of radially-orientated sandhoppers were pointed out (*G*-test, *G* = 6.784, df = 2, *P*<0.05, Fig. 6 C).

In tests of spectral filtering carried out with the vision of the sun prevented intact sandhoppers (Fig. 5J) and animals with the VP of their eyes occluded (Fig.5L) were mainly orientated in accordance with the expected direction. Instead, the mean orientation of individuals with the DP of their eyes black-painted were not concordant with their escape direction. Indeed, although the distribution obtained from these releases reaches the statistical significance (*V*-test, *u* = 2.295, *P*<0.05), most sandhoppers subjected to this treatment performed random directional choices and the mean resultant vector was deflected by 52° from the expected direction (Fig. 5 K).

The value of *v* measured from releases of animals with the upper edge of their eyes painted (*v* = 0.27) was lower than those registered from experiments carried out with controls (*v* = 0.67) and sandhoppers with the VP of their eyes obscured (*v* = 0.58). Moreover, comparisons pointed out that the distribution referring to releases of individuals with the DP of their eyes obscured differed statistically from that obtained in experiments conducted with intact animals (Fig. 5 J *vs* K, Watson’s *U*^*2*^ test, *U*^*2*^_(68, 34)_ = 0.347. *P*<0.005). Even significant differences in the frequencies of radially-orientated individuals were revealed (*G*-test, *G* = 15.888, df = 2, *P*<0.001, Fig. 6 D).

## DISCUSSION

Many terrestrial insects are known to rely on the skylight polarization pattern in their orientation (see Horvath and Varjù 2004, Wehner and Labhart 2006 for revisions). The existence of a specialized dorsal area in their compound eyes (dorsal rim area, DRA) is well proved, and several investigations demonstrated the involvement of ommatidia occurring in this region in the detection of the polarized light (see Wehner 1997, Labhart and Meyer, 1999, Horvath and Varjù, 2004; Wehner and Labhart, 2006 for reviews). The eyes of insects provided with the DRA also exhibit an eye regionalization of their visual pigments since UV photoreceptor cells are mainly located in the DRA in order to optimize the reliability of the celestial polarization pattern for their orientation behaviour, whereas those perceiving longer wavelengths occur in the rest of the eye (Waterman 1981, Wehner 1997, Cronin et al., 2014 Horváth and Varjù, 2004, Wehner, 1992, Rossel, 1989). Recently, studies of molecular phylogenetic have reported that insects and crustaceans are closely related, and this clade is called as Pancrustacea (Telford and Thomas, 1995; Friedrich and Tautz 1995; Pennisi, 2015) or Tetraconata, referring to the square ommatidia of many of its members (Dohle, 2001), suggesting the ancestor of the compound eye of this clade is same. Comparing with the huge number of research concerning with the regionalization of the compound eye, there are no investigation on crustacean eyes.

Our investigations conducted on the crustacean *T. saltator* revealed that there are morphological differences between DP and VP of the eyes. The pioneering research carried out by Bertolazzi (1937) on some species of gammarid amphipods (not *T. saltator*) show the existence of the fifth cell, however, Ercolini (1965) showed that the ommatidium of the compound eye of *T. saltator* possesses four retinular cells, but we found it possesses five retinular cells producing a square rhabdom. Four main rhabdomeres produced by R1-R4 were observed in all ommatidial in a compound eye, however the smaller R5 located in between R1 and R4 existed at DP, and the area of R5 rhabdomere decreased from median part to VP. The microvilli of R1 and R3 are oriented parallel to each other and orthogonally to those of R2 and R4, suggesting that photoreceptors within each ommatidium have differential sensitivities with respect to the direction of light polarisation. If Ercolini observed mainly the ventral part of the eye, he might not be able to find the rhabdomere of R5 which are mainly located in dorsal part. Our new morphological results indicate there are morphological regionalization, especially the localization of R5 depending on the eye region in the compound eye of *T. saltator*.

We assumed that the structural specializations of the retina determines the functional differences between DP and VP, therefore, we estimated roughly the visual filed from the light microscopic data. Since the number of the ommatidia of *T. saltator* increases with the growth of the body as reported for *Parhyale hawaiensis* (Ramos et al., 2019) and smaller ommatidial crystalline cones are frequently found at the peripheral region of the eye, we omitted these small ommatidia which possess small crystalline cones when we estimated the visual field. The visual field size in *T. saltator* were investigated using their pseudopupil (Beugnon et al., 1986); the individuals could potentially detect visual stimuli at 101 ° of vertical elevation. Our optical estimation of vertical elevation from the crystalline cone direction was wider, ca. 145°. Further experiment will be needed to confirm the origin of the visual field of *T. saltator*.

As expected (Cohen et al. 2010), both the waveforms of corneal negative- and positive responses were observed, however our data obtained from a proximal area were not stable showing both negative- and positive-responses. Therefore, we only used the corneal positive waveform obtained the electrode placed on the surface layer of crystalline cone (just below the cornea). There were no apparent differences in V-log I curves obtained from DP and VP, but the spectral sensitivity curve obtained from DP and VP showed the difference in shorter and longer peaks. The spectral sensitivity curve obtained from DP showed two distinct maxima, one in the long-wave range peaking at 550 nm and one in the UV range peaking at 430 nm, whereas the curves of VP possessed high sensitive region only around UV-blue and was decreased gradually in longer wavelength. Therefore, we can conclude there is a regionalization in the spectral sensitivity curves.

However, the relation between the electrophysiological data and morphological regionalization difference such as the localization of R5 is still unknown. Further studies such as intracellular recordings in different part of the compound eye are needed to solve this problem.

In the recordings from the dorsal rim area (DRA) in insect, the average spectral sensitivity curve showed a prominent UV responses (Waterman 1981, Wehner 1997, Cronin et al., 2014, Horváth and Varjù, 2004, Wehner, 1992, Rossel, 1989). These insects exploit skylight polarization for orientation or cruising-course control detecting the polarized sky light mediated by the ommatidia of DRA. Our morphological investigations revealed that the microvillar orientation of the pair R1 and R3 is orthogonal to the pair R2 and R4, expected to be sensitive to the polarisation of light. Our previous studies also showed the capability of sandhoppers to perceive polarized light in the blue band (λ = 435 nm) using a binary choice experiment while they do not use the skylight polarization pattern as a compass cue (Ugolini et al., 2013);therefore, the role of this factor in their celestial orientation has not been clarified in *T. saltator*. It has been speculated to be responsible for facilitating the detection/use of other cues by enhancing the intensity/spectral contrast across the sky (Ugolini et al., 2013). The broad spectral sensitivity curve obtained from DP showed two distinct peaks at blue and green region indicate it possesses at least two different receptor types. The necessary basic equipment for colour vison is to have at least two types of photoreceptors that differ in spectral sensitivity locating in approximately the same area (Skorupski and Chittka, 2011), therefore DP is thought to have colour vision. In insects, dung beetles also can use spectral cues for orientation. Testing them with monochromatic (green and UV) light spots in an indoor arena, the results suggest it could potentially use the celestial chromatic gradient as a reference for orientation (el Jundi et al., 2015). Our data strongly suggest the high possibility that DP retina of *T. saltator* possesses colour discrimination ability for orientation behaviour (Ugolini et al., 2013).

From the previous research, it is known that the visual pigments of animals have a maximum peak of α-band and a smaller peak of β-band exists at the shorter wavelength (Stavenga et al., 1993). The spectral sensitivity curves of VP obtained by ERG method show high sensitive region around UV-blue and was decreased gradually in longer wavelength. ERG recordings represent the overall spectral sensitivity of several ommatidia and therefore the summation of many retinular cells which were irradiated from a certain direction. It is difficult to find the peak in longer wavelength region in the spectral sensitivity curve of VP, however, there must be the retinular cell peaking at longer wavelength region. Because of those reason, the VP region also might have the colour discrimination ability.

Indeed, sandhoppers with the DP of their eyes painted met higher difficulties in orientating than the other groups of animals since in any experimental condition lower percentages of radially-orientated individuals were recorded and are more dispersed than intact individuals and animals with the VP of the eyes occluded. Moreover, in experiments of spectral filtering with the vision of the sun prevented their mean orientation was also deflected by 55° with respect that recorded for intact sandhoppers. In releases carried out with the vision of the sun precluded, the goodness of orientation of individuals with the upper area of their eyes painted reached fairly lower values than those recorded in releases of the other groups of individuals. Therefore, our investigations suggest that in *T. saltator* the dorsal 1/3 part is involved in the correct identification of celestial cues when the sun not visible. Instead, the rest of the eye seems not involved in the recognition of celestial orienting factors by this species. In fact, sandhoppers with the VP of their eyes obscured orientated in each experimental condition significantly toward their expected direction and frequencies of not radially-orientated lower than 40% were registered. Moreover, the goodness of orientation measured in releases of animals subjected to this experimental treatment reached in each case high values and in tests conducted under the natural sky and with the sun screened it even far exceeded the value obtained for control sandhoppers. However, it is important to underline that the goodness of orientation is influenced by the number of radially-orientated animals and its value increases as the number of individuals decreases (Batschelet, 1981). Since in releases under the natural sky and the vision of the sun prevented only 9 individuals subjected to the obscuring of the 2/3 VP of their eyes performed directional choices whereas 35 intact radially-orientated individuals were registered, the higher goodness of orientation value recorded for sandhoppers with the 2/3 VP area painted than controls (0.87 *vs* 0.56) was putatively due to the different sample sizes.

Therefore, our paper suggests that even *T. saltator* exhibits a regionalization of their visual capabilities necessary for celestial compass orientation (in the DP of its eyes) as well as many insects that use the pattern of polarized light as a compass cue (Waterman 1981, Wehner 1997, Cronin et al., 2014, Horváth and Varjù, 2004, Wehner, 1992, Rossel, 1989), *T. saltator* does not use the polarization pattern as a compass cue, but uses the intensity/spectral gradient across the sky (Ugolini et al., 2013). In fact, although a structural and functional eye regionalization has been pointed out in the stomatopod *Odontodactyllus scyllarus* (see Cronin et al., 2014), no evidence has never been provided on the involvement of specific eye regions in the orienting behaviour of crustaceans. Our investigations on *T. saltator* indicate for the first time in Crustacea the existence of an area located in the DP of its eye specialised in detecting certain wavelength ranges involved in the perception of skylight cues. We still don’t know the role of R5 dominant in DP, however, in *Parhyale hawaiensis*, the spectral absorption bands estimated from the genetic results differed between R1-4 and R5 (Ramos et al., 2019). Therefore, R5 cell of *T. saltator* also might have different colour absorption band. The ability of colour discrimination seems to make it easier to use the intensity and spectral contrast across the sky than the information of the polarization pattern. However, it is still possible to assume that *T. saltator* uses its polarization sensitivity (Ugolini et al., 2013) for their orientation. Both in DP and VP, the microvillar orientation of R1 and R3 are parallel, and that of R4 and R6 are also parallel, and those parallel microvillar orientation is orthogonal between the two pairs. Therefore, we can speculate that the VP uses the polarization of the reflected light from the sea water surface as a cue to the sea.

Studies of molecular genetics show that insects and crustaceans have evolved from a common ancestor (Dohle, 2001), therefore compound eyes are also thought to have common characteristics. In this paper, we show the regionalization, concerning with morphology, spectral response and celestial compass orientation, in the eye of a crustacean. However, there are several differences in the regionalization between the already reported insects’ vision and our findings in *T. saltator*. One of the most relevant is that in insects the part devoted to the perception of shorter wavelengths is the dorsal one. It is also well known that in the short wavelengths are perceived important celestial compass cues (e.g. see Horvath and Varjù 2004, Wehner and Labhart 2006, Dacke et al, 2011; Cronin et al. 2014). Despite sandhoppers use celestial cues in their orientation, shorter wavelengths are perceived by the lower part of the eyes. It seems quite strange, but we should underline that this regionalization is in agreement with the findings on the use of a coloured landscape in sandhoppers’ zonal recovery (Ugolini et al 2006). Tests made with artificial landscape (blue and green hemicycles) demonstrate a constant (compass) direction towards the blue hemicycle even when the directional indication coming from the sun compass was opposite to that of the landscape.

## Acknowledgements

We thank the Ente Parco Regionale di Migliarino, San Rossore, Massaciuccoli (Pisa) for the authorization to collect sandhoppers. Many thanks are due to Dr. Sofia Frappi for her help during experiments. We thank Preeminent Medical Photonics Educations & Research Center of Hamamatsu University School of Medicine for letting us use its equipment.

